# SARS-CoV-2 vaccine-breakthrough infections (VBIs) by Omicron (B.1.1.529) variant and consequences in structural and functional impact

**DOI:** 10.1101/2022.12.12.520021

**Authors:** Zainularifeen Abduljaleel, Sami Melebari, Saied Dehlawi, S Udhaya Kumar, Syed A. Aziz, Anas Ibrahim Dannoun, Shaheer M. Malik, C George Priya Doss

**Affiliations:** Science and Technology Unit, Umm Al-Qura University, P.O. Box 715, Makkah 21955, Kingdom of Saudi Arabia; Department of Medical Genetics, Faculty of Medicine, Umm Al-Qura University, P.O. Box 715, Makkah 21955, Kingdom of Saudi Arabia; Department of Molecular Biology, The Regional Laboratory, Ministry of Health (MOH), Makkah, Kingdom of Saudi Arabia; Laboratory of Integrative Genomics, Department of Integrative Biology, School of Bio Sciences and Technology, Vellore Institute of Technology (VIT), Vellore 632014, Tamil Nadu, India; Department of Pathology and Lab Medicine, University of Ottawa, 451 Smyth Road, Ottawa, ON, K1H 8M5, Canada; Department of Chemistry, Faculty of Applied Sciences, Umm Al-Qura University, Makkah, Kingdom of Saudi Arabia

**Author notes:** **Corresponding author:** Zainularifeen Abduljaleel, Ph.D. Research Scientist, Faculty of Medicine, Department of Medical Genetics, Umm Al-Qura University, P.O. Box 715, Makkah 21955, Saudi Arabia.

**Keywords:** SARS-CoV-2, Omicron (B.1.1.529), Delta breakthrough vaccine, Neutralizing antibody

## Abstract

This study investigated the efficacy of existing vaccinations against hospitalization and infection due to the Omicron variant of COVID-19, particularly for those who received two doses of Moderna or Pfizer vaccines and one dose of a vaccine by Johnson & Johnson or who were vaccinated more than five months previously. A total of 36 variants in Omicron’s spike protein, targeted by all three vaccinations, have made antibodies less effective at neutralizing the virus. Genotyping of SARS-CoV-2 viral sequencing revealed clinically significant variants such as E484K in three genetic mutations (T95I, D614G, and del142-144). One woman displayed two of these mutations, indicating a potential risk of infection following successful immunization, as recently reported by Hacisuleyman (2021). We examined the effects of mutations on domains (NID, RBM, and SD2) found at the interfaces of spike domains Omicron B.1.1529, Delta/B.1.1529, Alpha/B.1.1.7, VUM B.1.526, B.1.575.2, and B.1.1214 (formerly VOI Iota). We tested the affinity of Omicron for hACE2 and found that the wild and mutant spike proteins were using atomistic molecular dynamics simulations. According to binding free energies calculated during mutagenesis, hACE2 bound Omicron spike more strongly than SARS-CoV-2 wild strain. T95I, D614G, and E484K are three substitutions that significantly contribute to the RBD, corresponding to hACE2 binding energies and a doubling of Omicron spike proteins’ electrostatic potential. Omicron appears to bind hACE2 with greater affinity, increasing its infectivity and transmissibility. The spike virus was designed to strengthen antibody immune evasion through binding while boosting receptor binding by enhancing IgG and IgM antibodies that stimulate human *β*-cell, as opposed to the wild strain, which has more vital stimulation of both antibodies.

## Introduction

Globally, the Omicron variant of COVID-19 has spread rapidly and has been the most common variant in many countries since March 1, 2022. Omicron is far more contagious than other COVID-19 virus variants, such as Delta. The difference is modest deletions and insertions in two cases in the Omicron spike protein, which contains a minimum of 30 conservative amino acid substitutions (AAS). An AAS is detected in 15 cases in the receptor-binding domain (RBD) of the virus itself [1]. Several deletions and substitutions have affected genes related to other genomic regions. Delta Omicron appears to be more contagious than prior Omicron variants such as T478K, E484A, Q493R, G496S, Q498R, and key substitutions in spike protein of RBD E484A, G446S, G496S, G339D, S371L, S373P, S375F, S477N, N501Y, K417N, N440K, T478K, Q493R, Q498R, and Y505H, were significant since it has performed better than Delta as South Africa’s most prevalent variant [2]. There is evidence that it is linked to an increased risk of reinfection [3]. It also appears that Omicron replicates more rapidly than the Delta variant (World Health Organization). On November 24, 2021, in the South African epidemiological situation, Omicron B.1.1.529 T95I and D614G variants were initially reported to WHO [4]. The Delta variant was primarily responsible for the most recent peak. Even after successful vaccination and subsequent infection with a variant virus, there is still the possibility of illness in vaccinated individuals. Recent research indicates that mutations in the SARS-CoV-2 spike-RBD could affect how the virus connects with the host protein (ACE2). Like SARS-CoVs before it, SARS-CoV-2 enables spread by binding to ACE2 [5]. Thus, variants with a greater binding affinity for ACE2 suggest that they are most easily transmissible [6, 7]. In many Delta VOC sublineages and the Omicron VOC, T95I is an NTD mutation.

Randomized studies of convalescent plasma samples found that E484K reduced susceptibility by between 3 and 10 fold in 30% of models and by more than tenfold in roughly 10% of them [8–12]. D614G (SD2) prevalence started to increase in late February 2020, and within a few months, it had reached epidemic proportions. It has achieved a global rate of nearly 100% [13]. D614G disrupts contact between two or more promoters, increasing the likelihood of one or more of the three RBDs being open rather than closed [6, 14]. D614G is also thought to increase the number of spike proteins per virion [15, 16] and the S1/S2 cleavage [16, 17]. In some studies [14, 18], D614G viruses were more susceptible to mAb neutralization. Others were marginally more resistant to convalescent plasma and plasma from immunized people [14, 18].

Clustering is becoming one of the most popular tools for analyzing conformational ensembles [19]. Based on our first study, we investigated how mutations (D614G, T95I, E484K, and del142-144) affected the affinity and conformation of the spike in the WT strain with Delta/B.1.1.529, Alpha/B.1.1.7, VUM B.1.526, B.1.575.2, and B.1.1214 (VOI Iota) at the spike protein interface were found in domains (NID, RBM, and SD2). To better understand the risks posed by an individual or combine these mutations in the “second-wave” variants, we leveraged our *in-silico* simulation abilities to analyze WT and MT computationally. This will allow us to take novel computational methods to the next level and radically change the landscape of drug discovery.

## Materials and methods

### Collection of dataset

Spike SARS2’s FASTA sequence was obtained from the UniProt database (P0DTC2) using the reference sequence NP 828851.1. Based on Ezgi Hacisuleyman (2021), we gathered the nsSNPs from the Ensembl and dbSNP databases [20, 21]. The dbSNP database was used to extract functional annotations for each nsSNP. The Protein Data Bank (PDB codes: 6VXX and 6lzg) was used to obtain the structures of the Spike SARS2 protein [22, 23].

### Disease-causing non-synonymous SNPs (nsSNPs)

Using pathogenicity prediction tools, we predicted damaging or deleterious nsSNPs in the Spike SARS2 gene NCBI Reference Sequence: NC 045512.2. The support vector machines (SVMs) based PhD-SNP tool differentiates diseased mutations according to the nearby vicinity of substitutions [24]. An accurate consensus classifier for disease-related mutation prediction by PredictSNP [25] resulted in significantly accurate prediction and results for our mutations, confirming that consensus prediction is an accurate and robust method. SIFT [26], PolyPhen 2.0 [27], MAPP [28], and SNAP [29] web servers were used. They predict nsSNPs as neutral or disease based on a SIFT prediction score (destructive if the score is less than 0.05, neutral if the score is more significant than 0.05). PolyPhen assesses the effect of mutations as well as their phenotypic consequences on protein. In a previous communication, we explained the methodology for predicting deleterious nsSNPs [30, 31].

### Prediction of protein-destabilizing nsSNPs

The Gibbs free energy (*G*) change during protein folding represents protein stability. We used seven stability prediction tools in this study: SVib [32], ENCoM [33], ΔΔG mCSM [34], ΔΔG SDM2 [35], SAAFEC SEQ [36], DUET [37], and DynaMut [38]. The DynaMut program was used to perform this analysis. DynaMut is an approach for determining the effect of mutations on protein stability and uses a unified computational algorithm. The ΔΔG SDM2 algorithm distinguishes amino acid residue competence between WT and mutant proteins. To identify the variations in stability, SAAFEC SEQ, SVib, ENCoM, ΔΔG mCSM, and DUET use protein structural context and torsion angle prospects. The output includes information on the protein’s structural features and mutation sites, as well as the overall change in protein stability.

### Thermodynamic analysis of aggregation propensity

SODA is the latest algorithm for aggregation computation [39]. It forecasts changes in protein based on a variety of physicochemical characteristics. Protein-sol [40], PASTA 2.0v [41], and Aggrescan3D [42] are three other algorithms that predict aggregation-prone regions and inter-molecular pairings with *β*-strands for multiple protein sequences, with the results given as a function of sensitivity and specificity, as well as protein aggregation. Furthermore, using both programs together allows the calculation of differences in solubility of the protein based on the analysis of the tendency of protein sequences to have secondary structure, aggregation, hydrophobicity, and intrinsic disorder [39].

### Structure refinement to increase accuracy and immunogenicity analysis

A Spike variant with likely clinical importance includes one mutation (E484K) and three other mutations (D614G, T95I, and del142-144). We studied the structural effect of the D614G mutation on the S1/S2 interface of the domains. As described in PDB 6VXX, the chains labeled as A and B, corresponding to the S1 and S2 domains. ACE2 D614G substitution PDB: 6LZG-B of full-length human ACE2 was further analyzed by considering the enhancer promoter interactions (EPI) structure of S1 and S2 domains. The amber force field was used to construct the complex after introducing the appropriate RBD mutation E484K. The wild-type (WT) structures were determined by the 3Drefine refinement protocol involving atomic-level energy minimization and composite physics [43]. A rapid protein structure refinement was carried out on the optimized model using iterative hydrogen bonding network optimization and knowledge-based force fields from another MOE program (Molecular Operation Environment). The WT was evidently able to use mutagenesis embedded in the MOE to create D614G, T95I, and E484K mutations. We implemented the MOE algorithm to reduce the total potential energy of mutant and WT structures of a spike by releasing internal constraints and changing coordinate geometries. We also refined *in-silico* immunogenicity and immune response profiles via immunological simulations using the C-ImmSim algorithm [44].

### Molecular dynamics simulations

The crystal structure of the SARS-CoV-2 spike glycoprotein (closed state) with A Chain (6VXX-A) and molecular dynamics simulations of the spike complexed with ACE2 (6lzg-B) were investigated in this study using GROMACS [45]. Using the force field named GROMOS 54A7 [46], protein parameters were generated. We used the gmx genion, gmx solvate, and gmx editconf tools to build the triclinic simulation boxes 6LZG-B and 6VXX-A, with the first measuring 7.49×7.49×7.49 and the second 8.52×12.74×16.86. We used the simple point charge (SPC) model to solve the system and electro-neutralized it with the gmx editconf and gmx solvate tools. After applying the electro-neutralizing approach, we adjusted the structure to remove steric clashes and reduce energy using a solvate module for all systems. The systems were then immersed in a box with SPC16. The ions such as Na^+^ and Cl^-^ (0.15 M) contributed even more to the strategies for neutralizing and maintaining physiologic levels. All systems were minimized using the steepest descent of 1500 steps. The next step was a two-step equilibration. After heating the system to 300 K, we used NVT equilibration to stabilize its temperature within 100 ps. The system’s pressure and density were stabilized through the constant-pressure, constant-temperature (NPT) ensemble’s 100 ps. The final simulation of each structure resulting from the NPT equilibration phase lasted 100 ns. Finally, we examined the trajectory using several GROMACS analytic techniques. For time evolution of trajectory calculations, simulations were executed for 100 ns with a leapfrog integrator.

## Results

### Coordinate of Omicron variants and effective immune escape

A large number of mutations characterized the Omicron B.1.1.529 variants, including 26 to 32 changes in the spike (S) protein that is also found in three other variants of concern and are believed to make the virus more contagious, including c.1127C>T, p.(T95I), c.1841G>A, p.(D614G), and 1450G>A, p.(E484K) (Table 1). These three mutations occur in areas known to be involved in immune evasion. This region binds to the ACE2 receptor on the surface of human epithelial cells, resulting in virus membrane fusion and entry into the cell. This region also affects the ACE2 interaction surface, enabling the virus to evade the immune system and spread more widely. Omicron’s epitopes are located in the antibody-recognition areas for all three mutations. Interestingly, the proportions of SARS-Cov-2 variants have changed over time (Fig. 1a). Hence, monitoring variants has become a more significant concern. As a result of an interagency assessment of the variant’s attributes and public health risk in the Global Assessment (Fig. 1b), the classification was changed. According to the findings, the global number of confirmed 21K Omicron variant cases has increased significantly (Figs. 1c and 1d). Phylogenetic analysis revealed that the three Omicron families (21M Omicron, 21L Omicron, and 21K Omicron) are closely related (Fig. 1e).

**Table 1.**
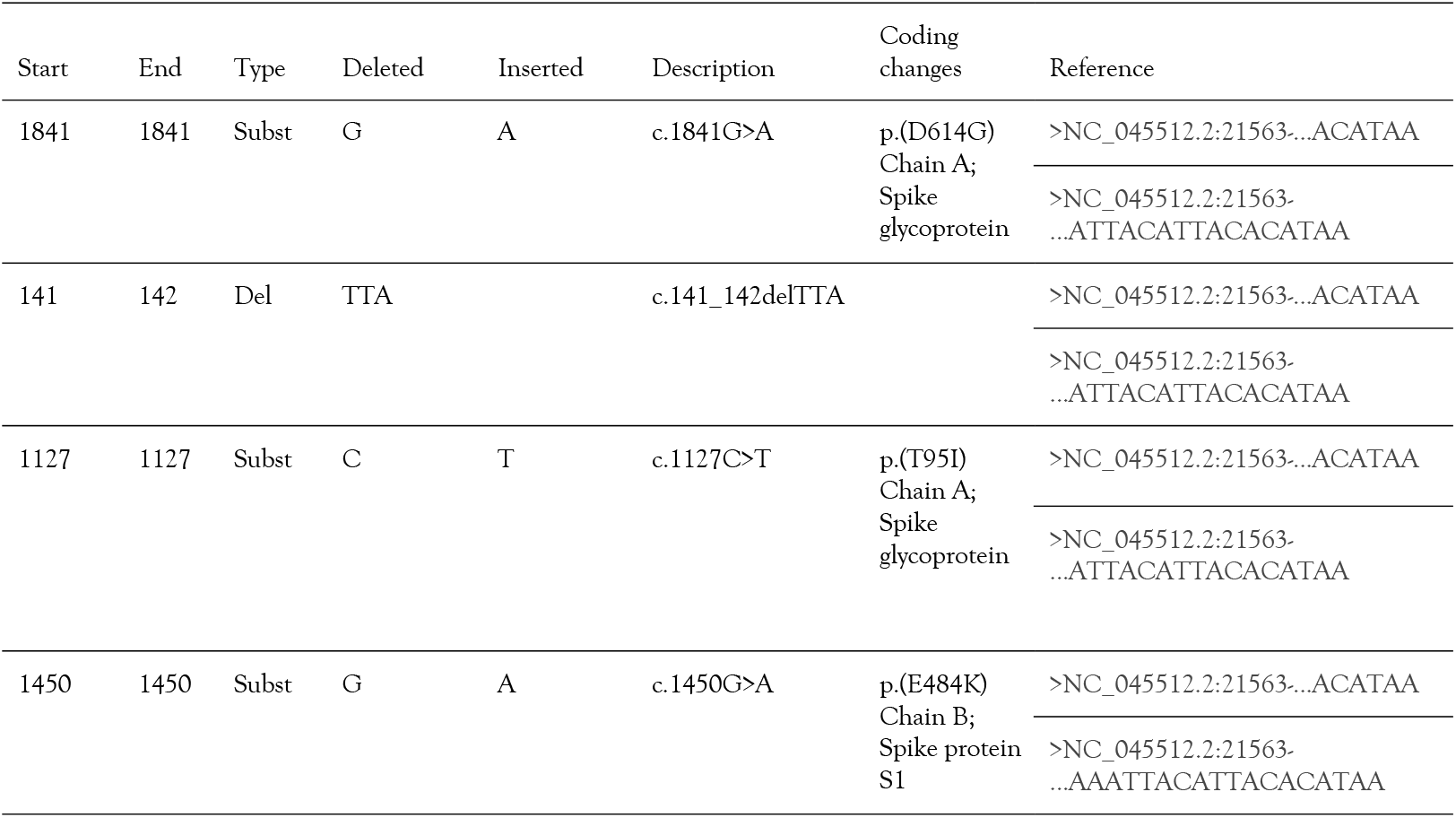
Genome coordinate of omicron immune escape mutations

**Figure 1.**
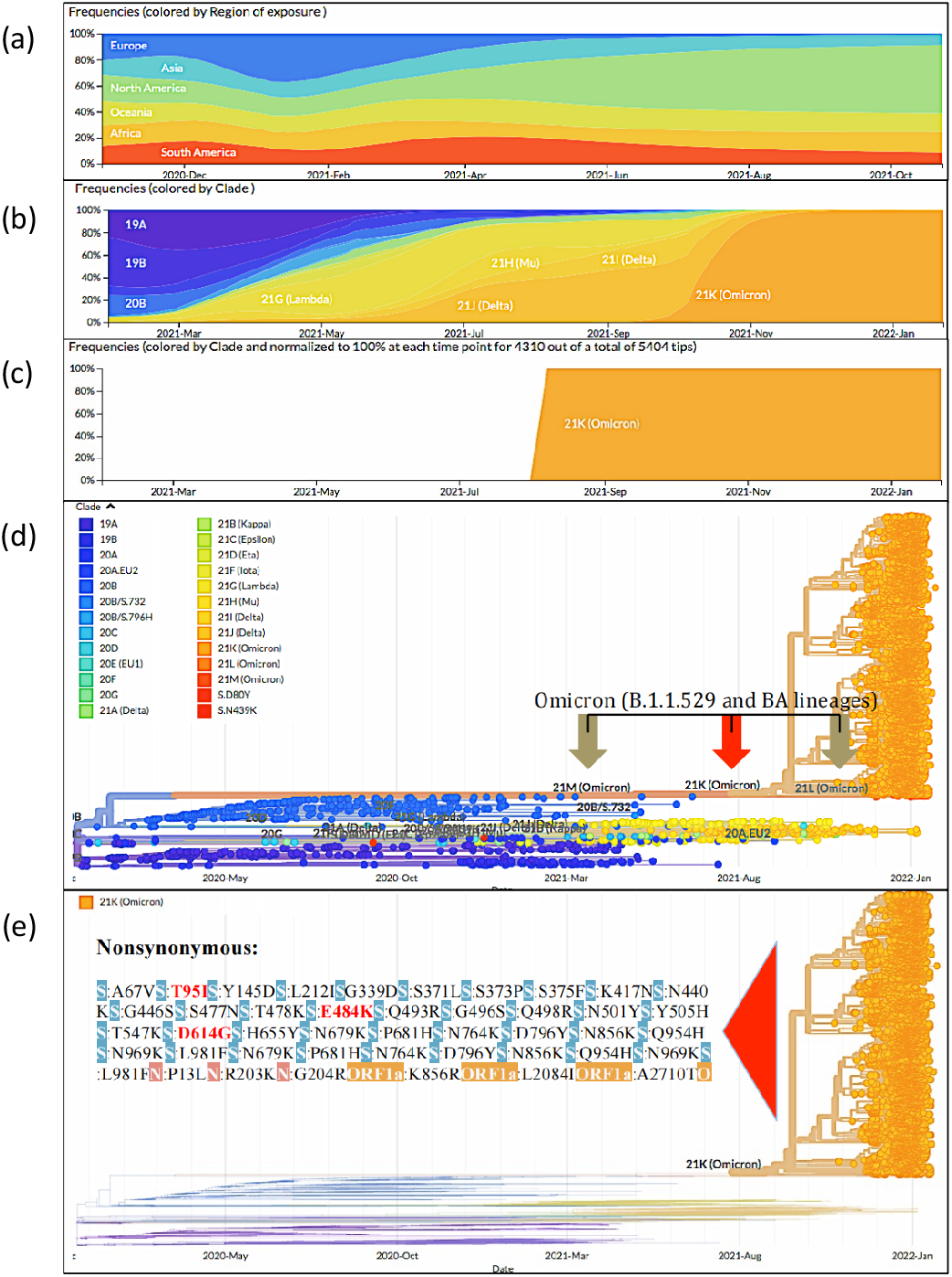
(a) Frequencies encoded in different colors representing different regions. (b) Different SARS-CoV-2 variants have different frequencies encoded in different colors. (c) Frequencies colored by clade and normalized to 100% at each time point for 4310 SARS-CoV-2 Omicron variants out of a total of 5404. (d) Phylogenic tree displaying a variety of Omicron variants (B.1.1.529) in 21M and 21L Omicron-highlighted color (green), 21K Omicron-highlighted color (red). (e) 21K Omicron non-synonymous variants are organized in different colors, with variants that escape immune response highlighted in red (T95I, D614G, and E484K).

### Prediction of deleterious and non-deleterious SNPs

The three nsSNPs chosen (T95I, D614G, and E484K) were assigned to seven different programs: SIFT, SNAP, PhD-SNP, MAPP, PolyPhen-2, PolyPhen, and PredictSNP. However only five were predicted to be damaged: PredictSNP (61%), PolyPhen-1 (74%), PolyPhen-2 (68%), SIFT (79%), and SNAP (72%). T95I (P0DTC2) mutations in strains B.1.1.529 (Omicron), B.1.526 (Iota), and B.1.1.318 were mapped from UniProt P0DTC2 position 95. Other strains with D614G mutations are B.1.1.529 (Omicron), B.1.1.7 (Alpha), B.1.351 (Beta), B.1.429 (Epsilon), B.1.526 (Iota), B.1.1.318, P1, P2, 20A.EU1, and 20A.EU2. When grown in primary human epithelial cells, EU2 binds to hACE2 more effectively, producing more infectious particles. It does not affect the neutralizing properties of SARS-CoV-2. However, pseudotyped vesicular stomatitis virus (VSV) particle production increased ex vivo due to mutagenesis. This strain produces more infectious particles when it is cultured in primary human epithelial cells, enhances replication and transmissibility, and is not deleterious E484K (P0DTC2). We sent the query sequence for the nsSNPs to the webserver in FASTA format. SIFT analysis projected nsSNPs to cause 79% of the threshold at which discrimination is significantly damaged (score=0.05), and also predicted nsSNPs to have tolerable (score>0.05) impacts on the spike protein. Out of three nsSNPs, Polyphen-2 predicted T95I to be “possibly damaged”. Based on the SIFT and PolyPhen-2 findings, we selected SNPs with a PolyPhen score of >0.90 and SIFT of 0.05 to enhance prediction accuracy. The results of both SIFT and PolyPhen v2 improved prediction accuracy by excluding SNPs with SIFT scores of 0.05 and PolyPhen scores of less than >0.90. Table 2 summarizes the results achieved using these techniques. These SNPs could then be investigated using other methods that were likely to be detrimental.

**Table 2.**
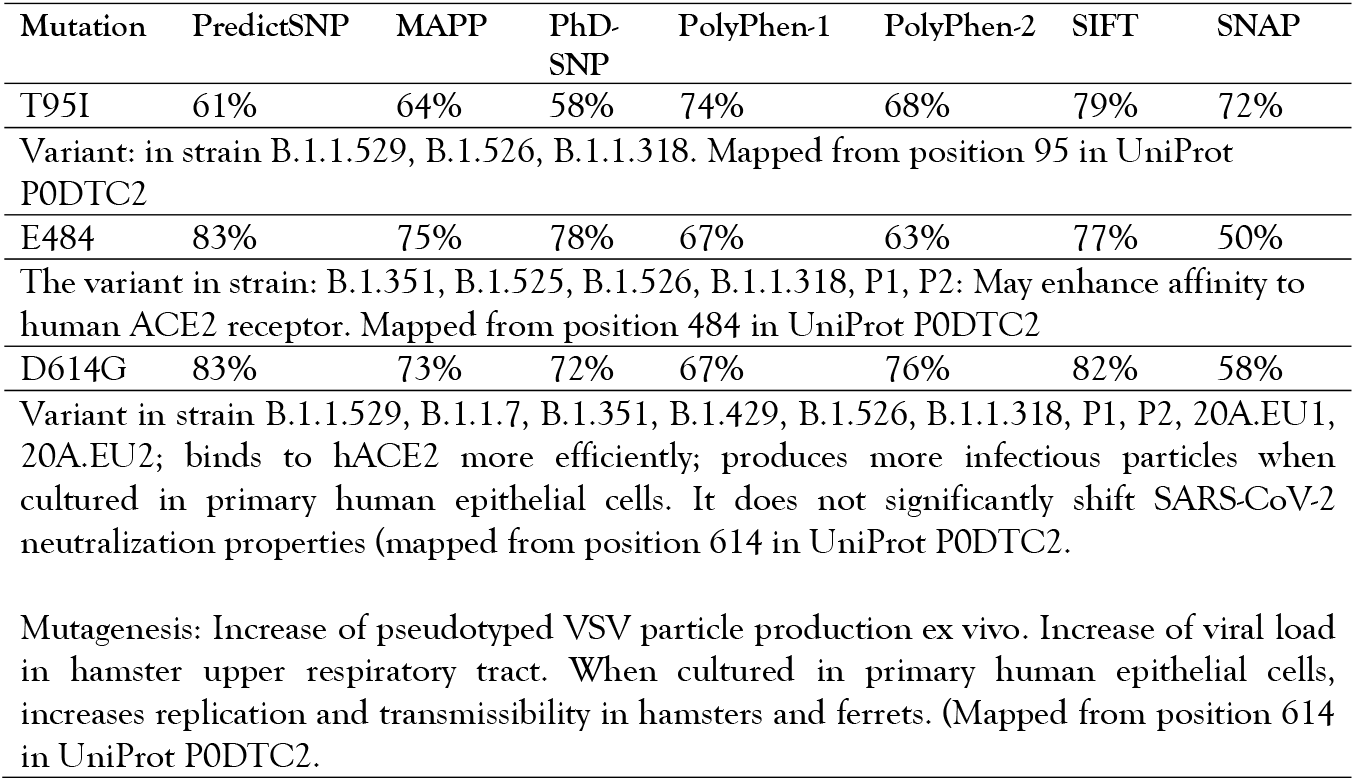
Accurate consensus classifier for prediction of disease-related mutationsm

| Mutation | PredictSNP | MAPP | PhD-SNP | PolyPhen-1 | PolyPhen-2 | SIFT | SNAP |
| --- | --- | --- | --- | --- | --- | --- | --- |
| T95I | 61% | 64% | 58% | 74% | 68% | 79% | 72% |
| Variant: in strain B.1.1.529, B.1.526, B.1.1.318. Mapped from position 95 in UniProt P0DTC2 |  |  |  |  |  |  |  |
| E484 | 83% | 75% | 78% | 67% | 63% | 77% | 50% |
| The variant in strain: B.1.351, B.1.525, B.1.526, B.1.1.318, P1, P2: May enhance affinity to human ACE2 receptor. Mapped from position 484 in UniProt P0DTC2 |  |  |  |  |  |  |  |
| D614G | 83% | 73% | 72% | 67% | 76% | 82% | 58% |
| Variant in strain B.1.1.529, B.1.1.7, B.1.351, B.1.429, B.1.526, B.1.1.318, P1, P2, 20A.EU1, 20A.EU2; binds to hACE2 more efficiently; produces more infectious particles when cultured in primary human epithelial cells. It does not significantly shift SARS-CoV-2 neutralization properties (mapped from position 614 in UniProt P0DTC2). |  |  |  |  |  |  |  |
| Mutagenesis: Increase of pseudotyped VSV particle production ex vivo. Increase of viral load in hamster upper respiratory tract. When cultured in primary human epithelial cells, increases replication and transmissibility in hamsters and ferrets. (Mapped from position 614 in UniProt P0DTC2). |  |  |  |  |  |  |  |

### Stability, flexibility, and conformational dynamic changes in structures

With the help of MD results, we analyzed the effects of T95I, D614G, and E484K mutations on the structural integrity of spike proteins. Except for the loop in the side chain of the two mutations D614G and E484, after 100 ns of simulation, they became more flexible; and in WT, no major structural alterations were observed. In the case of T95I, the C-terminal helix structure was observed to be lost. The RMSD calculated was used to investigate the overall changes in spike protein stability caused by mutations. There are five different algorithms for predicting changes in protein thermodynamic equilibrium stability due to point mutations. These computational methods elucidate the differences in backbone flexibility between mutant and WT atoms. ΔΔSVib ENCoM −4.531 kcal/(mol·K) was calculated as the average of standard mode analysis of energy between WT and MT and values of T95I WT (decrease of molecule flexibility). Although the two algorithms were classified as destabilizing, the difference between them for ΔΔG mCSM was only 0.103 kcal/mol whereas for SAAFEC SEQ it was only 0.33 kcal/mol (again, destabilizing).

In comparison, the other two mutations were indicated as stabilizing by two algorithms, ΔΔG SDM 1.010 kcal/mol (stabilizing) and ΔΔG DUET 0.353 kcal/mol (stabilizing), as shown in Table 3. Overall, the thermodynamic stabilization values of two mutants (D614G and E484K) did not differ significantly, and there were no significant variations between the two simulation runs in any of the structure parameters. These findings also revealed no major structural alterations in the probability distribution function values of the WT and mutants.

**Table 3.** Thermodynamic stability changes upon point mutation

| Normal mode analysis energy between wild-type and mutant |  |  |  |  |  |
| --- | --- | --- | --- | --- | --- |
| Mutation | $\Delta\Delta S_{\text{vib}}$ ENCoM | $\Delta\Delta G$ mCSM | $\Delta\Delta G$ SDM | SAAFEC SEQ | $\Delta\Delta G$ DUET |
| p.(D614G) | −4.531<br>kcal/(mol·K)<br>(decrease of<br>molecule<br>flexibility) | −0.425 kcal/mol<br>(destabilizing) | 1.180 kcal/mol<br>(stabilizing) | −1.02 kcal/mol<br>(destabilizing) | 0.079 kcal/mol<br>(stabilizing) |
| p.(T95I) | −4.646<br>kcal/(mol·K)<br>(decrease of<br>molecule<br>flexibility) | −0.103 kcal/mol<br>(destabilizing) | 1.010 kcal/mol<br>(stabilizing) | −0.33 kcal/mol<br>(destabilizing) | 0.353 kcal/mol<br>(stabilizing) |
| p.(E484K) | 0.171 kcal/(mol·K)<br>(increase of<br>molecule<br>flexibility) | −0.080 kcal/mol<br>(destabilizing) | −0.630 kcal/mol<br>(destabilizing) | −0.08 kcal/mol<br>(destabilizing) | 0.128 kcal/mol<br>(stabilizing) |

### Structural and functional impact of Omicron variants

About 50 genetic mutations are present in the Omicron coronavirus, 36 of which occur at the spike protein. However, three variants, 6LZG-B chain E484K, 6VXX-A chain D614G, and T95I, fall within an antigenic supersite region. We evaluated the effects of the three mutations by the following criteria: interactions established by the modified residue, structural domains within which the altered residue was located, and alterations to known and mutant forms of the residue. The short indicated that wild-type and Omicron S protein variants had distinct structural variations (Figs. 2a–2d). The first mutation transformed threonine to isoleucine at position 95. The hydrophobicity of the mutant 95I residue was higher than that of the wild-type T95 residue. According to UniProt, the wild-type residue was engaged in a cysteine bridge, crucial for protein stability. Cysteines can only form these connections; the mutation eliminates this contact, which deleteriously influences the protein (Fig. 3a). Because of the loss of the cysteine bond and this residue, the structure has destabilized. UniProt revealed that T95I provides an amino acid, identified in the same domain as BetaCoV S1-NTD, with unique characteristics that can disrupt and eliminate this domain’s function.

**Figure 2.**
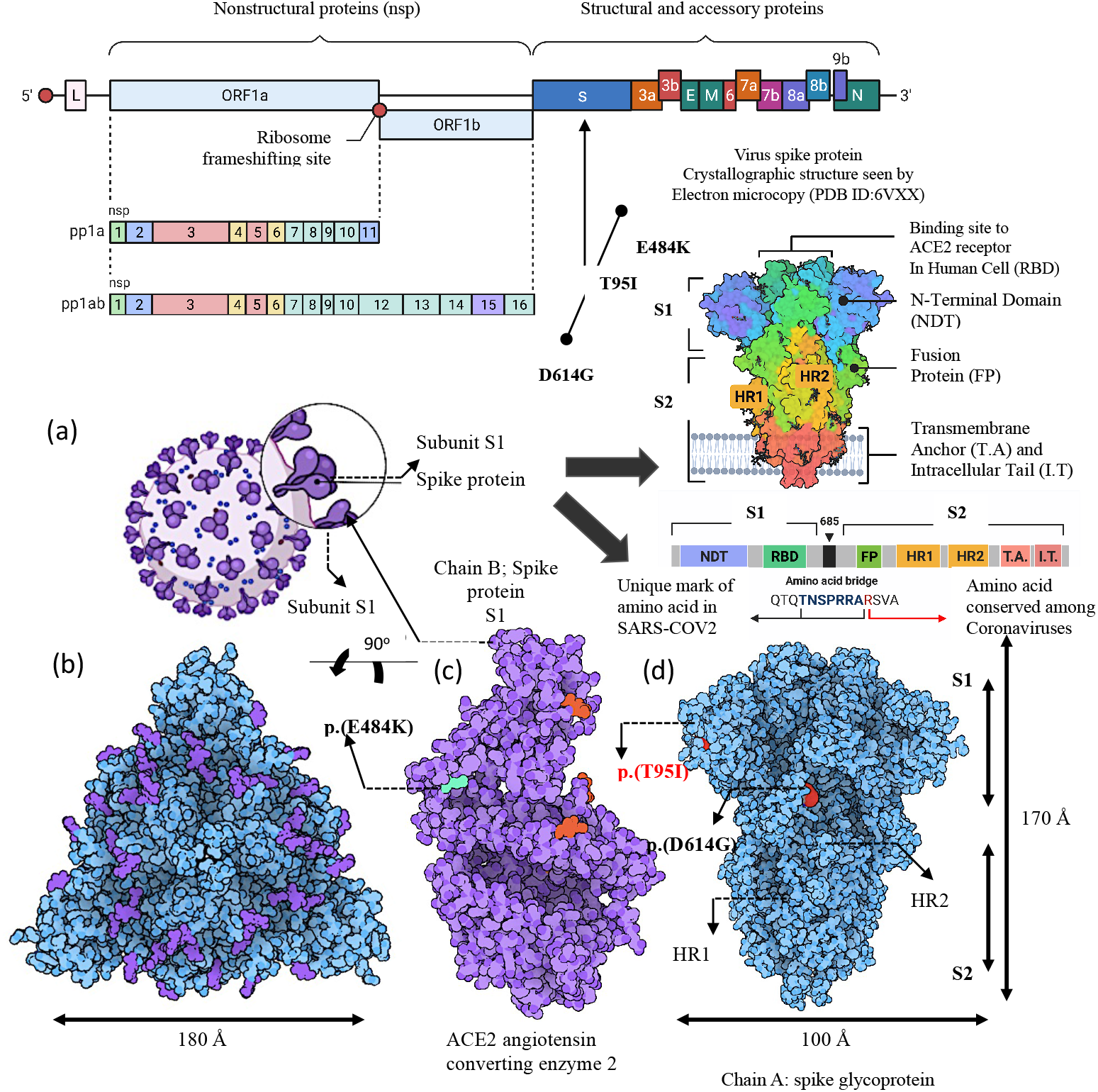
Genomic structure organization of SARS-CoV-2. (a) Based on the SARS-CoV-2 Wuhan-Hu-1, the representation of the SARS-CoV-2 genomic structure. The genome is divided into two domains: non-structural and accessory proteins. The S protein is made up of two subunits (S1 and S2) which are separated by the S cleavage site. Open reading frame (ORF), spike (S), envelope (E), membrane (M), nucleocapsid (N), N-terminal domain (NTD), receptor-binding domain (RBD), fusion peptide (FP), heptad repeat 1, and heptad repeat 2 (HR1 & 2), contain the core binding motif in the external subdomain, transmembrane anchor (TA), and intracellular tail (IT). (b, d) Spike protein in open form with two substitution mutations (RCSB Protein Data Bank ID 6VXX-A chain) p.(T95I) and p.(D614G) highlighted in a trimer axis vertical view (left) and an orthogonal top-down view along this axis (right). (c) A close-up view of the receptor-binding domain (RBD) bound to hACE2 with identified mutations (RCSB Protein Data Bank ID 6VXX-A chain) and RBD residues p.(E484K) shown as molecular spheres colored by amino acid variant frequency and hACE2 depicted in red. Positions near the RBD–hACE2 interface have a high frequency of amino acid variants. 1 □=10^-10^ m

**Figure 3.**
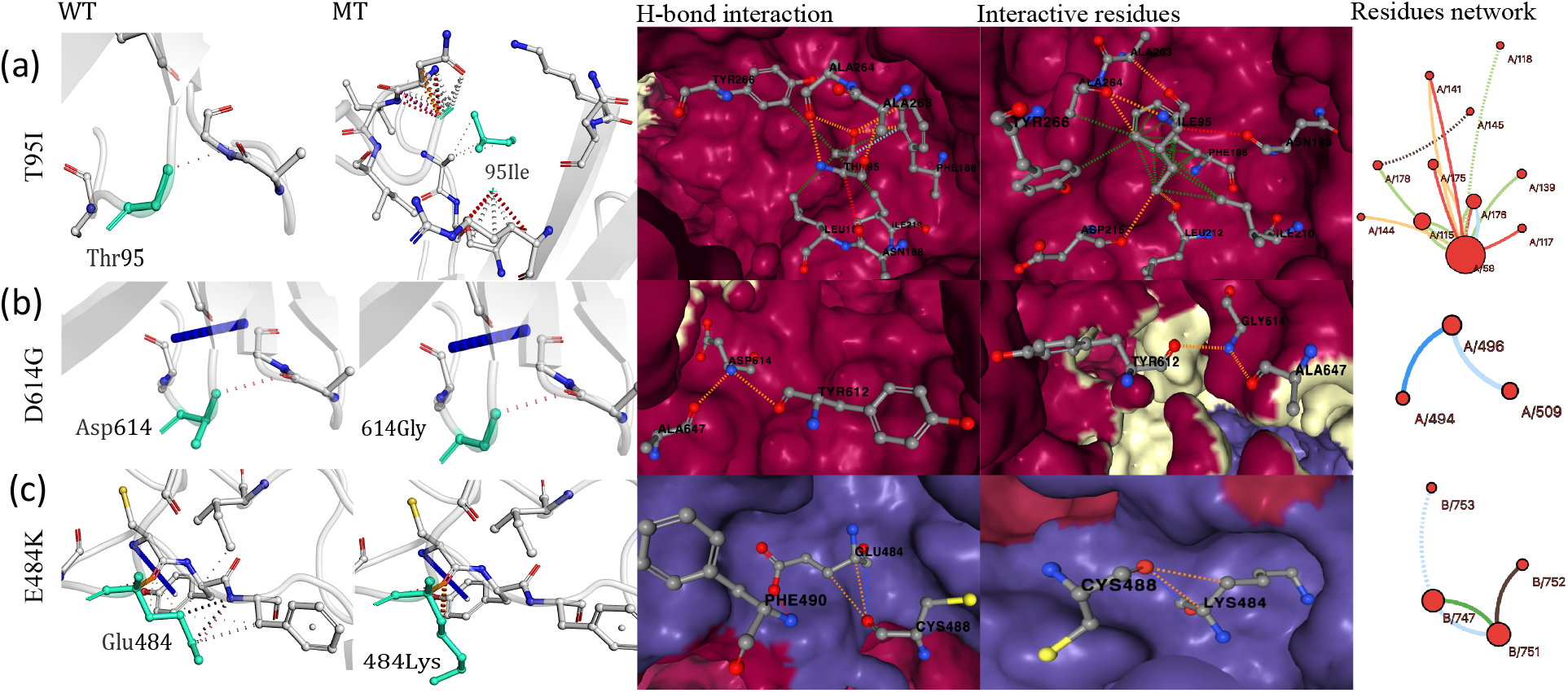
Location and distribution of wild and mutant residues in the SARS-CoV-2 protein. Three unique spike monomers form a complex to form the spike protein. (a) The mutation T95I at spike protein might significantly affect the spike protein structure. S protein residue T95I at the cleavage site affects fusion, facilitation, and efficient entry into host cells. Wild-type SARS-CoV-2 residue T95 is represented in red and mutated residue 95I in yellow. The mutated residue is marked in contact atoms. H-bond interacting residues (ALA263, ALA264, ILE95, PHE186, ASN188, ILE220, LEU212, ASP215, TYR266, and ALA264) were found in the PDB: 6VXX chain A/141, A/118, A/145, A/117, A/176, A/158, A/144 and A/178 and substantially strengthen the hydrogen bond while replacing T95I. (b) The structural mutation in the SARS-CoV-2 spike protein occurs at position 614. The wild type of ASP614 is highlighted in red, while the receptor-binding region is depicted in purple. Intermolecular atomic contacts between 614 in a reference spike and two residues (ALA647 and TYR612) in its adjacent spike-protein monomer chain. These two contact residues have a hydrophilic to hydrophobic repulsive action and are unstable, which is abolished in the D614G mutation when aspartate is replaced by glycine (Table 2). Only three interacting residues were found in the PDB: 6VXX chain A/496, A/509, and A/494. H-bond interaction was not substantially stronger because hydrogen bonds were lost as a result of the D614G mutation. (c) Another rendering of E484K’s position on RBD is in purple. The RBD of S protein interacts hydrophilically with the N-terminal helix of the hACE2 receptor. One residue (CYS488) strongly interacted with hACE2 and RBD (PDB: 6lzg_chain B). Other intact intermolecular residues were B/751, B/753, B/752, and B/484. H-bond interaction was not substantially stronger because hydrogen bonds were lost due to the E484K mutation.

Despite the fact that the wild-type residue was very conserved, there were a few different types at this location. There were no mutant residues or other residue types with similar attributes in other homologous sequences at this location. Based on the conservation scores, this mutation was most likely detrimental to the protein. The mutant residue was near a highly conserved site and introduced a more hydrophobic residue at this location. It caused hydrogen bonding to break and correct folding in a state of disruption. The 6VXX (A) 95 THR > ILE: (Buried H-bond breakage) substitution impeded all side-chains with side-chain H-bond(s) and side-chains with main-chain bond(s) created by a buried THR residue (RSA 0.7%). This substitution impeded all side chains with side-chain H-bond(s) and side-chains with a main-chain bond(s) created by a buried THR residue.

At position 614, the second aspartic acid to glycine mutation occurred. Each amino acid was distinct in size, charge, and hydrophobicity. The original wild-type residue D614 and the newly inserted mutant residue 614G displayed many differences in these features. The mutant residue was about half the size of its wild-type counterpart and had a neutral charge, while the wild-type residue had negative control. The mutant residue 614G had higher hydrophobicity than the wild-type residue D614. The mutation added glycine to this position. Glycine is a highly flexible amino acid that can cause the protein’s essential stiffness to become disrupted at this point (Fig. 3b). In other homologous sequences, this mutant residue was discovered at this place more frequently, indicating that the mutant residue occurred in more proteins than the wild-type residue. The D614G mutation in question is unlikely to be detrimental to the protein. This residue was adjacent to a highly conserved location. The size of wild-type and mutant amino acids also differed. Interactions were lost because the mutated residue was smaller. At this position, the mutation created more hydrophobic connections and induced hydrogen bonding to break, causing the correct folding to be disrupted.

The size of the third mutant residue, 484K, was more extensive than that of the wild-type residue. The mutant residue exhibited a positive charge in contrast to the wild-type residue’s negative charge. According to UniProt, the wild-type residue was engaged in a cysteine bridge, crucial for protein stability. Cysteines can only form these connections; the mutation eliminates this interaction, strongly influencing the protein’s 3D structure (Fig. 3c). The discrepancies between the old and new residues created structural instability in combination with the loss of the cysteine bond. The mutation introduced a unique amino acid that disrupted and eliminated the function of the BetaCoV S1-CTD domain. The mutation occurred in a stretch of residues that UniProt has classified as the RBD. Differences in amino acid characteristics can disrupt the function of this area. Compared to other homologous sequences, there were neither mutant residues nor residues with similar properties at this site. Based on the conservation scores, this mutation was most likely detrimental to the protein. The mutation added a residue with the opposite charge to that of the wild type. Other protein residues or ligands may be repellent to these residues.

### Immunogenicity of SARS-CoV-2 Omicron S protein variants

According to immunoinformatics, the amino acid sequence of the S protein from UniProt ID P0DTC2 evoked an immune response in both wild and Omicron strains. Despite having comparable immunological features, the immune response outcomes of the wild type and Omicron were significantly different. The immunological response was shorter in the three Omicron variations (T95I, D614G, and E484K). Antibody titers were higher for the Omicron version, which contained IgM+IgG, IgG1+IgG2, and IgG1 alone due to immunogenicity (Fig. 4). Despite the fact that the wild type had more active B-lymphocytes than the mutant, they were similar in class-II presentation, internalized Ag, anergic response, and reproduction. In the S protein wild type, the number of T-regulatory memory lymphocytes was more numerous, while the number of T-regulatory non-memory lymphocytes was smaller. Omicron activity and resting achieved 89 (cells per mm^3^), but the wild type only reached 80 (cells per mm^3^). The number of regulatory T-cells (but not memory cells) was higher in Omicron. The S protein promoted the same number of cytokines and interleukins in both cell types. TGF-b and IFN-g were the most common immune-system stimulators for both forms of S protein. IFN-γ levels were 620,000 ng/mL in the wild type and 630,000 ng/mL in Omicron.

**Figure 4.**
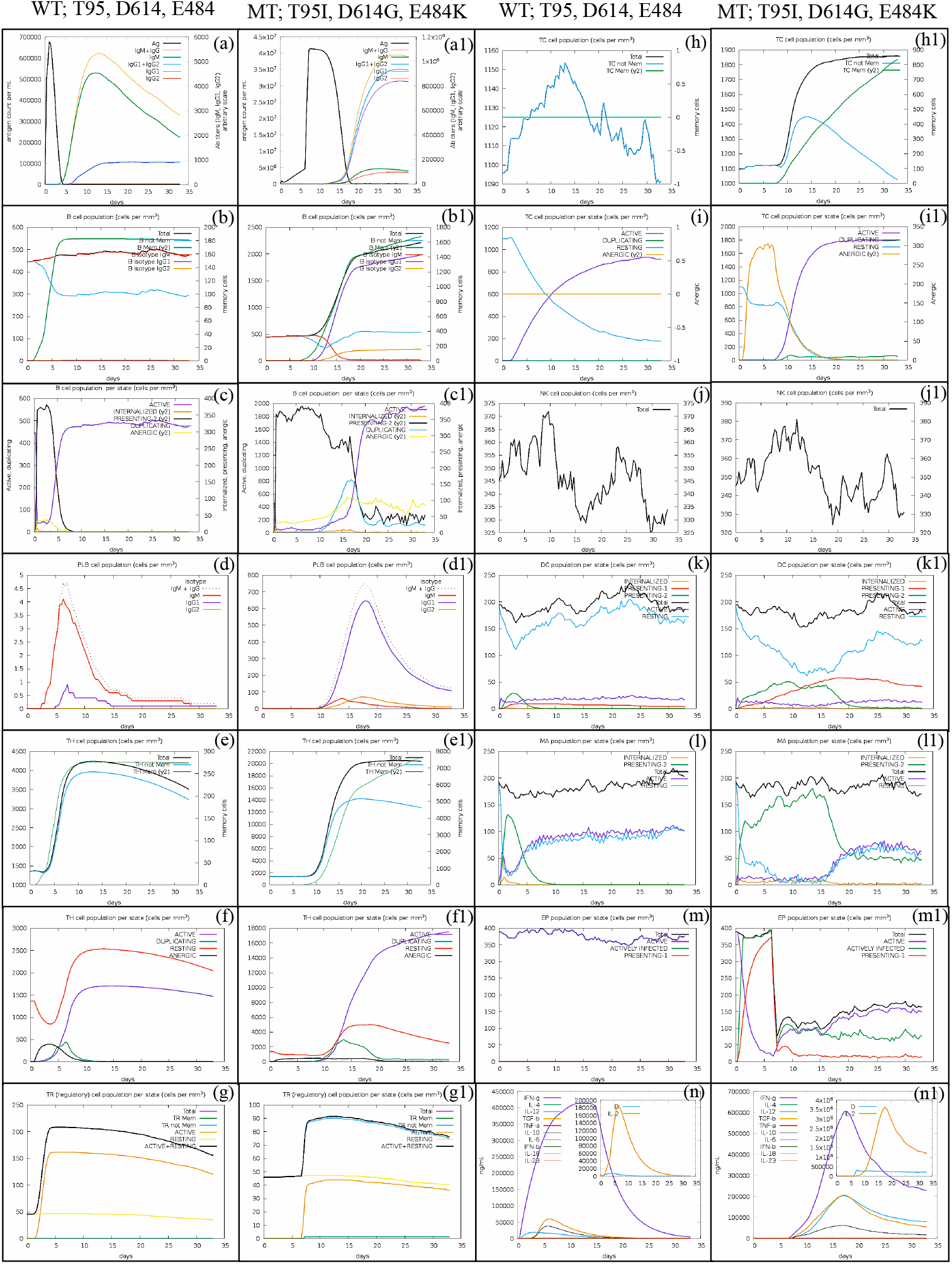
Immune response to Omicron S proteins that are wild-type and mutant. (a, a1) Ag count per mL and all Ab titers as measured after 30 days of infection. Peaks of IgG1+IgG2, IgM+IgG, and IgG1 are higher in Omicron than in wild type. (b, b1) and (c, c1) Population of B lymphocytes per entity state, showing counts for active, presenting on class II, internalizing the Ag, duplicating, and anergic lymphocytes. The presenting class II number in Omicron is higher, but the counting activity is less than in the wild type. (d, d1) PLB cell population. (e, e1) and (f, f1) TH cell population. (g, g1) CD4 T-regulatory lymphocyte count. Plots show the total, memory, and individual entity-state counts. The activity and resting of Omicron reach 89 (cells per mm^3^) while it reaches 80 (cells per mm^3^) for the wild type. Omicron has a higher level of regulatory T-cells but not memory. (h, h1) and (I, i1) TC (T-cell) population. (j, j1) Natural killer (NK) cell population. (k, k1) Dendritic cell (DC) population. (l, l1) MA population. (m, m1) Electroporation (EP) population. (n, n1) The inset plot’s concentration of cytokines and interleukins is a warning sign. Both types of S protein stimulate similar levels of cytokines and interleukins. Among the types of S proteins, IFN-g and TGF-b are most strongly stimulated by the immune system, and IFN-γ of the wild-type is 620,000 ng/mL and 630,000 ng/mL for Omicron.

### Analysis of structural aggregation and residue interaction propensity

Using the four independent algorithms Protein-sol, SODA, Aggrescan3D, and PASTA in our pipeline, along with a unique spike protein sequence, we anticipated the aggregation potential of each mutant. The mutations did not affect the protein’s assembly ability in most cases. But in at least one of the employed methods, 40% of the mutant sequences increased the aggregation proneness score. To begin, each method’s output score for predicting the likelihood of protein aggregation was standardized to the score of the reference strain. The input sequences were considered to be at risk of aggregation if their AggreScore was greater than 0. Since the majority of the mutations found in these variations were in the S protein’s structural domains, structural instability could be related to an increase in the APR’s aggregation score. Most of the changes discovered in these variants were in the S protein’s structural domains, suggesting that the structural destabilization was attributable to an increase in aggregation. We measured the region under the curve for the S protein wild-type and three (T95I, D614G, and E484K) mutants of more significance ΔΔG (Figs. 5a–5i) to evaluate protein solubility and energy (Table 4). Compared to the protein’s wild-type counterpart, this curve indicated that several amino acids had higher aggregation scores. However, neither the peak area’s percentage nor its tendency to aggregate was significantly altered by our alterations. We further investigated the average contact maps of polar and non-polar groups of S-protein with close contact residues to gain more insight into the physical driving force between molecular interactions. To accomplish this, we compute the average interatomic distance between the atoms of the closed two residues (Figs. 6a–6c). The predicted interresidue contacts and distances from the SARS Cov-2 S protein were less than eight amino acids. Contact density was calculated for each protein by dividing the total number of non-local contacts (residue pair with seq_sep≥6).

**Figure 5.**
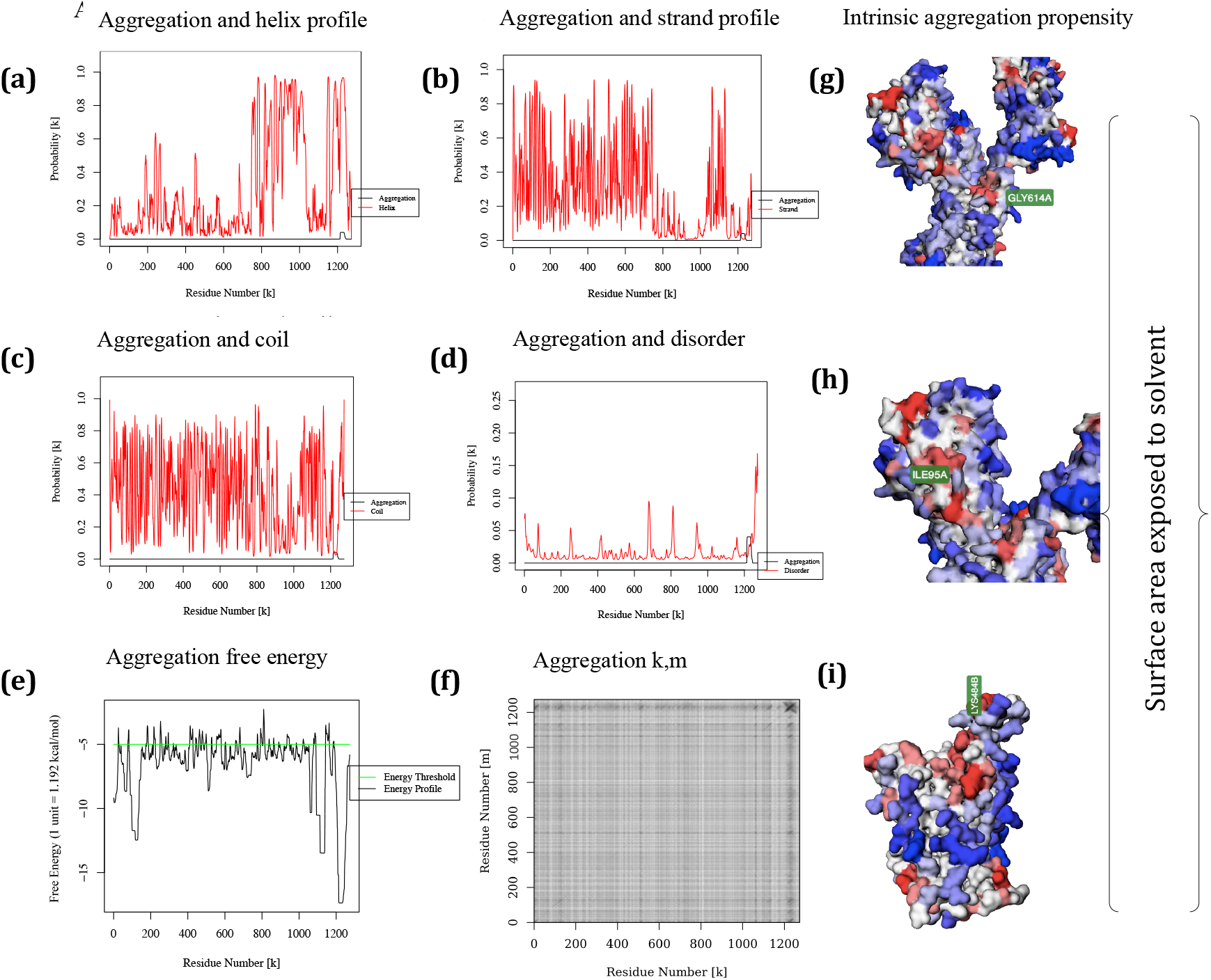
(a) Aggregation and helix profile. (b) Strand intrinsic aggregation propensity profile. (c) Aggregation and coil. (d) Aggregation and disorder. (e, f) Aggregation free energy. Intrinsic aggregating tendency residues are colored based on their A3D value, spanning from blue to red colors signifying aggregation-prone residues with solvent exposure. (g) Gly614 (h) Ile95 and (i) Lys484.

**Table 4:** Prediction of protein solubility from disorder and aggregation propensity

| Sequence | Helix | Strand | Aggregation | Energy disorder | SODA |  |
| --- | --- | --- | --- | --- | --- | --- |
| Wild type | 0.3 kcal/mol | 0.289 kcal/mol | -6.674 kcal/mol | 0.016 kcal/mol |  |  |
| T95I | -0.297 kcal/mol | 1.014 kcal/mol | -60.4 kcal/mol | 0.064 kcal/mol | -59.046 kcal/mol | Less soluble |
| D614G | -0.334 kcal/mol | 0.315 kcal/mol | 2.59 kcal/mol | 0.038 kcal/mol | 2.616 kcal/mol | More soluble |
| E484K | 1.263 kcal/mol | -1.174 kcal/mol | 0.196 kcal/mol | 0 | 0.495 kcal/mol | More soluble |

**Figure 6.**
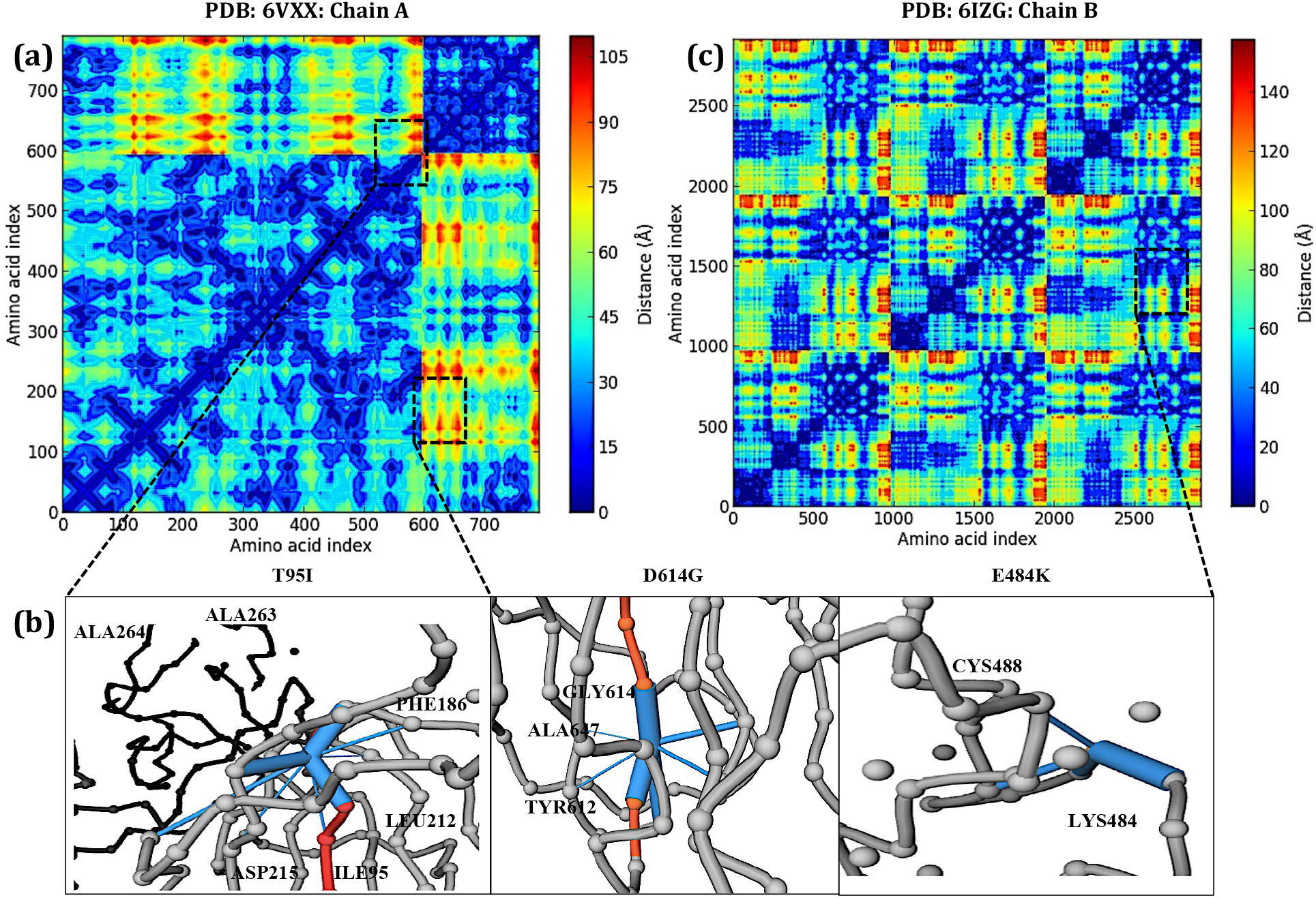
Putative signaling of protein residues and ligand contacts. (a, c) Large sets of feasible deep maps reproduce a near-native residue contact matrix using an 8 □ threshold. (b) The ligand-binding pocket of spike protein bound to the ligand contact color in blue; PDB 6LZG-B with chain B residue variants (T95I and D614G); 6VXX-A with chain A residue variant (E484K). All residues that come into direct contact with each other are displayed as grey sticks. The inter residues are colored to indicate the number of Cα atomic interactions and their structural properties. Contacts are defined as a function of distances up to 1 nm using the positions of all heavy atoms. The calculated Cα atom distances were in Ile96 (10 □), Gly614 (14 □), and 484 (11 □). 1 □=10^-10^ m

### Molecular dynamics (MD) simulation and root-mean-square deviation of atomic positions

We ran molecular dynamics simulations for 100 ns (using GROMACS) for the selected proteins 6LZG-B chain and 6VXX-A chain after generating two systems for the 6LZG-B chain (6LZG-E484K) and another 6VXX-A chain with two mutations (6VXX-D614G and T95I). This allowed us to distinguish the conformational behavior and stability differences between the proteins. As a result of the RMSD deviations, the protein’s stability of its conformation could be determined throughout the simulation. The less deviation there was, the more stable the protein structure. The RMSD values 6LZG and 6VXX are plotted against time in Figs. 7a and 7b. The RMSD of 6LZG wild protein was (0.308±0.045) nm on average, but the average 6LZG RMSD value of mutant E484K was (0.404±0.106) nm. The average RMSD values of mutant E484K were higher than those of wild-type, stating that they were less stable. The 6VXX wild-type’s standard RMSD was (1.547±0.289) nm. D614G and T95I mutants had average 6VXX RMSD values of (1.389±0.224) nm and (1.765±0.358) nm, respectively. T95I mutants had higher standard RMSD values than the 6VXX wild-type, which was less stable.

**Figure 7.**
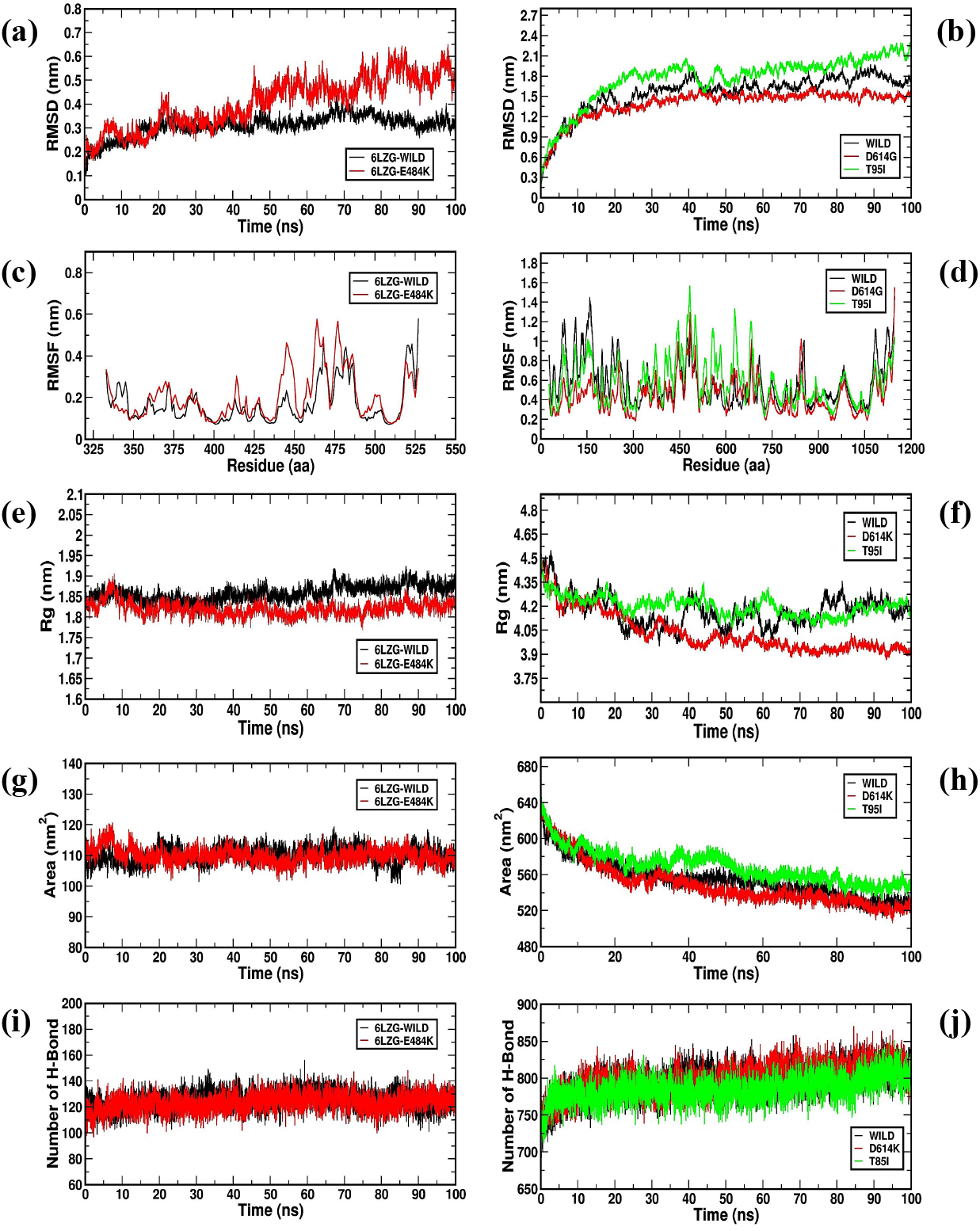
Analysis of RMSD, RMSF, Rg, SASA and the total number of hydrogen bonds of wild and mutation of 6LZG and SPIKE protein respectively. (a) Root-mean-square deviation (RMSD) of the Ca atoms in 6LZG wild and mutant. (b) Root-mean-square deviation (RMSD) of the Ca atoms in spike, wild and mutant. (c) RMSF values of the alpha carbon over the entire simulation of 6LZG wild and mutant. (d) RMSF values of the alpha carbon over the entire simulation of spike, wild and mutant. (e) Radius of gyration (*R*_g_) over the entire simulation of 6LZG wild and mutant. (f) Radius of gyration (*R*_g_) over the entire simulation of SPIKE, wild and mutant (g) Solvent accessible surface area (SASA) of 6LZG wild and mutant. (h) Solvent accessible surface area (SASA) of spike, wild and mutant (i) Total number of H-bond count throughout the simulation of 6LZG wild and mutant. (i) Total number of H-bond count throughout the simulation of SPIKE, wild and mutant.

### Root mean squared fluctuation of the S-protein monomers

RMSFs for the 6LZG and 6VXX proteins and all mutants were calculated using the gmx rmsf tool to predict their flexibility and stability. High RMSF values indicated that the structure was more flexible during molecular dynamics simulation (MDS). Following that, a plot displaying RMSF of mutants vs. 6LZG and 6VXX residues is shown in Figs. 7c and 7d. The RMSF of 6LZG protein was (0.171±0.096) nm on average. The average RMSF value of the mutants E484K was (0.205±0.112) nm. Also, we discovered that the E484K had greater values (RMSF) at several 6LZG binding sites, resulting in ligand-binding issues. The RMSF of the 6VXX was (0.565±0.232) nm on average. D614G and T95I mutants had average RMSF values of (0.444±0.181) nm and (0.576±0.228) nm, respectively. The T95I mutant also had ligand-binding issues resulting from greater RMSF at several 6LZG binding sites. We used the gmx gyrate tool to calculate the *R*_g_ values for the wild-type proteins 6LZG and 6VXX and all mutant proteins. Reduced *R*_g_ levels demonstrated the protein’s native form to be more compact and rigid. The *R*_g_ values were plotted against time for the respective proteins (Figs. 7e and 7f). The *R*_g_ value of the 6LZG was *1.9 nm on average. The average *R*_g_ value of the mutants E484K was *1.8 nm. The *R*_g_ value of the 6VXX wild-type protein was *4.05 nm on average. D614G and T95I mutants had average *R*_g_ values of *4.02 and *4.35 nm, respectively. These results show that all mutants were more compact and rigid than the wild-type protein.

### Solvent-accessible surface area (SASA) of WT and MT

The solvent-accessible surface area was calculated using the GMX SASA tool values for 6LZG, 6VZZ, and all mutant proteins. Figs. 7g and 7h depict a plot of the SASA values of wild-types and mutants versus time. The 6LZG wild-type protein had an average SASA value of *115 nm^2^. The average SASA value of the E484K mutants was *110 nm^2^. The E484K protein and T95I were found to have a greater SASA value than the 6LZG wild-type, showing that mutant proteins’ hydrophobic cores were more exposed to the solvent than those of the wild-type. The 6VXX wild-type protein had an average SASA value of *560 nm^2^. The D614G and T95I mutants had average SASA values of *550 and *575 nm^2^, respectively.

### Hydrogen bond analysis of wild and mutant proteins within the time span of the current simulation

Additional hydrogen bonds suggest a more stable protein structure because H-bonds are essential for its stability. The gmx hbond tool counted the number of H-bonds in the wild-type proteins 6LZG, 6VXX, and all mutant proteins. Then, we plotted these numbers against time, as shown in Figs. 7i and 7j. In the 6LZG wild-type protein, the median number of intramolecular H-bonds was *130. In the case of E484K mutants, the average number of H-bonds was comparable to that in 6LZG wild-type. Approximately, 820 was the median number of intramolecular H-bonds in the 6VXX wild-type, and the numbers in D614G and T95I mutants were comparable.

## Discussion

The global spread of Omicron (B.1.1.529) 21K is spreading much faster than Delta, infecting people who have already been vaccinated or have recovered from COVID-19 infection [47]. Given that the Omicron variant is rapidly displaced (Fig. 1), it is reasonable to assume that the clinical profile described represents a disease caused by the Omicron variant since Delta variants have been identified in the region [9]. The current COVID-19 vaccines target the S protein [48]. The S protein structure is essential for any variant, and obtaining this structure allows us to understand the Omicron variant better. The multiple RBD mutations involved in Omicron would affect an antibody and RBD complex, potentially resulting in multiple cancellations for each antibody and RBD complex leading to potential vaccine breakthrough over different mutations. Some of the mutations in Omicron are also found in other VOCs. They may be associated with immune escape, enhanced transmissibility via induction of cell fusion, and susceptibility to treatment, as demonstrated theoretically or through functional studies of previous variants [49, 50]. Six mutations in the N-terminal domain may be linked to antibody neutralization evasion by innate, vaccine-based, or monoclonal antibodies. According to a recent study, viral sequencing revealed potentially clinically significant variants such as E484K and three mutations in one woman (T95I, D614G, and del142–144). These findings point to a probable risk of sickness as a result of effective vaccination and later infection with a viral variant. They support ongoing efforts to prevent, diagnose, and characterize variants in vaccinated people [51]. These three mutations, T95I, D614G, and E484K, all occur within the NTD supersite and are likely to disrupt antibody binding as well [52]. They include Omicron B.1.1.529 (69–70) variants, also found in Alpha B.1.1.7, Beta B.1.351, Delta B.1.617.2, Gama P.1 (D614G and E484K), and T95I, which was also variants found in Kappa and Iota. Immune evasion ability and greater transmissibility have been linked to the receptor-binding domain and the furin cleavage site [53].

This study looked into the structural and functional implications of non-synonyms in S protein Omicron variants. The disease-causing pathogenic spectrum was screened using advanced computational methods. The strains identified in Wuhan, ID (6VXX Chain-A and 6LZG Chain-B) were used as a template reference from PDB. Finally, the atomic levels of three pathogenic mutations (T95I, D614G, and E484K) that cause distinct protein structure changes with immune response were determined using MD simulation. The SARS-CoV-2 variant with the spike D614G mutation spreads rapidly and becomes the dominant form of the virus. The D614G variant is linked to increased infectivity and viral load [13].

Another study revealed that the D614G mutation disrupted a salt bridge between D614 and K854, decreasing the S1 and S2 subunits [6]. The D614G mutation in the N-terminal domain (NTD) is not clinically pathogenic, according to the deleterious functional analysis (Table 1), and has no effect on the neutralizing characteristics of SARS-CoV-2. When the D614G mutation was thermodynamically analyzed using five different algorithms, only three were found to be destabilizing at a maximum of −4.531 kcal/(mol·K), meaning a decrease in molecular flexibility (Table 2). Furthermore, the aggregation propensity of the solubility result, 2.616 kcal/mol, was slightly different between wild and mutant forms (Table 3). Experimental results show that the E484K mutation, in particular, prevents the virus from detecting antibodies [54]. The functional analysis results clarify that the E484K mutation is not harmful compared to other algorithms. The E484K mutation had maximum thermodynamic stability of 0.171 kcal/(mol·K), indicating an increase in molecular flexibility. In addition, the disorder of aggregation propensity had a solubility of 0.495 kcal/mol. Based on the significant differences between the D614G and E484K mutations, E484K may allow the virus to escape antibodies. The non-polar, non-charged Thr residue is replaced with a non-polar, aliphatic Ile residue in the T95I mutation. As a result, these mutations alter the protein’s overall milieu in the replacement region. In the D138Y mutant, a bulkier aromatic Tyr residue replaces a shorter, negatively charged hydrophilic residue, creating a notable change in the surrounding environment. The polar acidic Asp residue replaces a Gly residue that lacks a side chain in the G142D mutant, increasing its ability to form hydrogen bonds.

Five out of seven functional analysis outcomes were predicted to be harmful. The thermodynamic stability analysis resulted in −4.646 kcal/(mol·K) (a decrease of molecular flexibility), whereas the tested solubility disorder of aggregation propensity resulted in −59.046 kcal/(mol·K), also indicating a decrease of molecular flexibility (Table 4). Immune stimulation for immunogenicity analysis determined T95I, D614, and E484K on strain S protein in all mutations compared to wild and mutant forms. As a result, strain S protein activated the immune system, producing a differentiated reaction, unlike wild strain, which stimulated IgG and IgM more strongly (Fig. 4).

In addition, we performed molecular dynamics simulations to analyze the change in structural behavior between the WT protein and the proteins with mutations. Plots were created for the protein with E484K mutation and analyzed against the WT in the protein with PDB ID 6lzg, and the D614K and T95I mutations were plotted against WT in the protein with PDB ID 6VXX. We analyzed the RMSD to check the convergence of the molecules over the simulation period [55]. The RMSD of the protein with mutation E484K exhibited a higher deviation than the WT (Fig. 7a). In contrast, the protein with D614G mutation showed a smaller variation than the WT, and T95I mutation showed a higher deviation than WT (Fig. 7b). However, the convergence of 100 ns simulation plots shows that the trajectory output of the molecules is suitable for further analysis. The RMSF was explored to understand the differences in the residual level fluctuation of the protein [56]. The RMSF of the protein with mutation E484K exhibited greater fluctuation than the WT (Fig. 7c). The RMSF of the proteins with mutation D614G showed less fluctuation than the WT, and the T95I mutation showed higher fluctuation than the WT (Fig. 7d). This reveals that the protein with mutations E484K and T95I is capable of influencing its own crucial residues of the protein and thereby impacting the binding pockets. We performed *R*_g_ analysis to understand the differences in compactness between WT and mutant proteins.. The WT and the protein with E484K showed stable deviation in the radius of gyration (Fig. 7e).

On the other hand, the WT protein with PDB ID 6VXX and the proteins with mutations D614K and T95I showed deviation from the beginning of the simulation to the end (Fig. 7f). The surface area of a biomolecule accessible to a solvent is known as solvent-accessible surface area (SASA) [57]. The WT protein and the protein with E484K mutation showed a similar area accessible to solvent (Fig. 7g). On the other hand, though the WT protein and protein with D614K mutation had similar SASAs, that of the protein with T95I was comparatively higher. This change in the SASA of T95I might influence the molecules’ free energy transfer due to mutation [58]. Finally, we calculated the difference in intramolecular H-bonds to understand the reason for changes in deviation, fluctuation, SASA, and other structural changes (Fig. 7h). The change in several hydrogen bonds determines the changing structure of the protein [59]. The WT and the protein with E484K showed a similar number of H-bonds throughout the simulation period (Fig. 7i). The number of intramolecular H-bonds was comparatively decreased in protein with the T95I mutation compared to the WT protein and protein with the D614K mutation (Fig. 7j). This decrease in intramolecular H-bonds due to the D614K mutation appears to be the reason for higher structural deviation, as well as changes in compactness, fluctuation, and SASA of the protein (Figs. 7a–7h). Compared to three Omicron mutations, T95I, D614, and E484K, two of the mutations, T95I and D614, had theoretically strong global outbreaks lineages of vaccine-breakthrough SARS-CoV-2 infections.

## Conclusions

In order to examine Omicron’s infection effects, we used a computational model which has been successfully trained and tested using several experimentation datasets. It has been shown that Omicron is ten times more contagious than the original virus and twice as infectious as the Delta variant, based on our study.

Based on the complex structures of antibody-RBDs, we find that Omicron is about twice as effective as Delta at escaping vaccinations. In order to protect against Omicron infection symptoms, a stronger vaccination is required. Immune evasion in healthy subjects who had received three doses, including booster doses, of different COVID-19 vaccines, illustrates the ineffectiveness of current immunizations against Omicron. Therefore, Omicron-targeted vaccines are essentially required to combat all SARS-CoV-2 strains.

## Abbreviations

VBIs: Vaccine-breakthrough infections
COVID-19: Coronavirus
ACE2: Angiotensin-Converting Enzyme 2
RBD: Receptor-binding domain
NTD: N-terminal domain
RBM: Receptor binding motif
SD2: Subdomain
AAS: Amino acid substitutions
WHO: World Health Organization
MD: Molecular dynamics
WT: Wild type
MT: Mutant
dbSNP: Single nucleotide polymorphism database
PDB: Protein data bank
NVT: Equilibrate the system
FASTA: Fast Alignment Search Tool
SIFT: Sorting intolerant from tolerant
SNAP: Screening for non-acceptable polymorphisms
PROVEAN: Protein variation effect analyzer
S1-CTD: Predictor of human deleterious single nucleotide polymorphisms (PhD-SNP) Subunits C-terminal domain
RMSD: Root-mean-square deviation
RMSF: Root mean square fluctuation
TGF-b: Transforming growth factor
IFN-g: Interferon-gamma
SODA: Protein solubility based on disorder and aggregation
PASTA: Prediction of amyloid structural aggregation
VOC: Variant of concern
SASA: Solvent-accessible surface area
PLP: Proteolipid protein
HR1 & 2: Heptad repeat 1 heptad repeat 2
TA: Transmembrane anchor
IT: Intracellular tail
MOE: Molecular operating environment
RSA: Relative solvent accessibility
DC: Dendritic cells
EP: Electroporation
NID: Immunomodulatory drugs
Rg: Radius of gyration
VSV: Vesicular stomatitis virus
EPI: Enhancer Promoter Interactions

## Acknowledgments

The authors would like to thank the Deanship of Scientific Research at Umm Al-Qura University for supporting this work by Grant Code (22UQU4340573DSR001).

## Author contributions

Conceptualization: ZA and GPDC; methodology and data analysis: ZA, SM, SD, UKS, SAA, AID, and SMM; writing original draft preparation: ZA, SM, GPDC, SD, and UKS; writing review and editing: ZA, SAA, and GPDC. All authors have read and agreed to the published version of the manuscript.

## Declarations

### Conflict of interest

The authors declare that they have no conflict of interest.

### Ethical approval

This article does not contain any studies with human or animal subjects performed by any of the authors.

### Availability of data and materials

All data analyzed during this study are included in this article.

### Disclaimer

The study sponsor had no involvement in the ‘study’s design, data collection, data analysis, data interpretation, or report writing. The first and corresponding authors had full access to all study data and had the final say on whether or not to submit the paper for publication.

### Collaborators

The authors would also like to thank the management of Vellore Institute of Technology (VIT), Vellore, India, and the Department of Pathology and Lab Medicine, University of Ottawa, for their assistance with the analysis of the research.

